# Cytotoxic T lymphocytes targeting a conserved SARS-CoV-2 spike epitope are efficient serial killers

**DOI:** 10.1101/2022.01.24.477535

**Authors:** Mohsen Fathi, Lindsey Charley, Laurence J.N. Cooper, Navin Varadarajan, Daniel Meyer

## Abstract

Understanding the cellular immune response to infections, cancers and vaccines lags behind the investigation of humoral responses. While neutralizing antibody responses wane over time, the ability of T cells to recognize viruses including SARS-CoV-2 is instrumental to providing long-term immunity. Although T-cell receptor (TCR) repertoire screening can provide insights into the skewing of a T-cell response elicited upon vaccination or infection, they unfortunately provide no assessment into the functional capacity of T cells or their ability to eliminate virally infected targets. We have used time-lapse imaging microscopy in nanowell grids (TIMING) to integrate the migration of individual T cells with analysis of effector functions including cytokine secretion and cytotoxicity. Machine learning is then applied to study thousands of videos of dynamic interactions as T cells with specificity for SARS-CoV-2 eliminate targets bearing spike protein as a surrogate for viral infection. Our data provide the first direct evidence that cytotoxic T lymphocytes from a convalescent patient targeting an epitope conserved across all known variants of concern (VoC) are serial killers capable of eliminating multiple infected targets. These data have implications for development of vaccines to provide broad and sustained cellular immunity and for the recovery and monitoring of individuals who have been exposed to SARS-CoV-2.

**Multidisciplinary abstract:** We present an imaging platform that uses artificial intelligence (AI) to track thousands of individual cell-cell interactions within nanowell arrays. We apply this platform to quantify how the T cell component of adaptive immunity responds to infections. Our results show that T cells specific for a conserved epitope within the SARS-CoV-2 spike protein are serial killers that can rapidly eliminate virally infected targets. The ability to map the functional capacity of T cells and their ability to kill infected cells provides fundamental insights into the immunology of vaccines and recovery from infections.

**Graphical Abstract:** 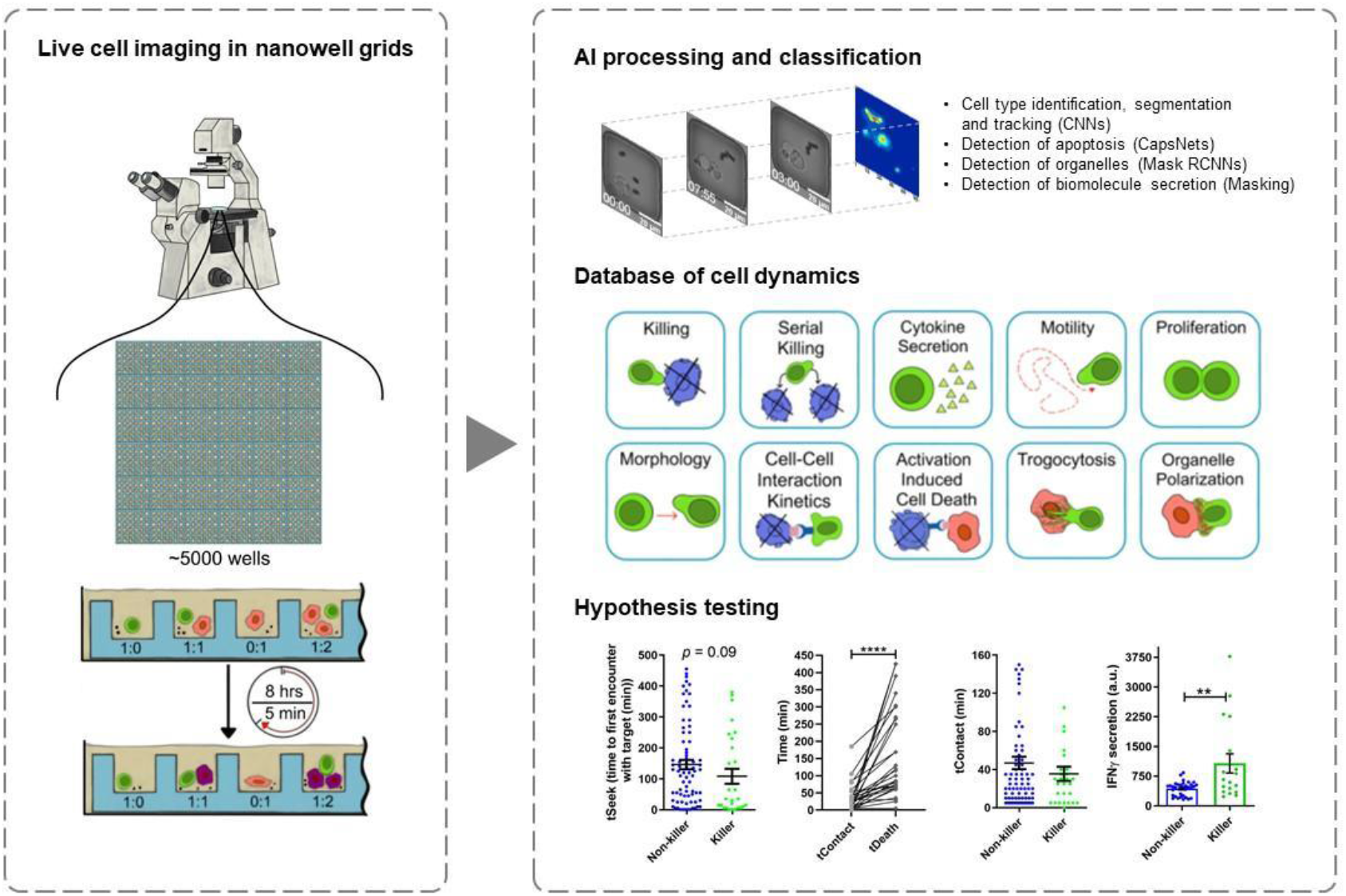

## Introduction

Commercial vaccines seek sterilizing immunity by generating neutralizing antibodies that can protect against viruses like SARS-CoV-2^1^. The immunity, however, wanes over time and erodes protection and exposes individuals to risk of disease and communities to outbreaks of infection^2^. The cellular component of adaptive immunity mediated by cytotoxic T lymphocytes (CTLs) plays a fundamental role in the recognition and removal of virally infected respiratory cells^3^. Understanding the functional potency of CTLs against SARS-CoV-2 is important in predicting ongoing cellular immunity^4^. Improving our understanding T-cell responses to COVID-19 will help develop next-generation vaccines with an ability to offer protection against emerging variants^5^. Furthermore, analysis of T-cell immunity will help characterize patients with asymptomatic through severe disease.

In response to infection and vaccination, the humoral immune response is typically well characterized, primarily because of the ease of studying antibody molecules in vitro^1^. Studying T-cell mediated cellular immune responses is much more challenging since the response is mediated by the cells and not the soluble TCR^6^. T cells are capable of a complex array of functions including migration to infected tissue, activation, killing of infected cells, secretion of cytokines/chemokines and proliferation^6^. Mapping all of these functions onto the same cell requires the ability to study the spatiotemporal changes in cellular behavior across thousands of cells.

## Assay principle

This application note describes the use of time-lapse imaging microscopy in nanowell grids (TIMING), an integrated microscopy and artificial intelligence (AI) platform that can image and profile the dynamics of thousands of cells and cell-cell interactions simultaneously^7^. The properties measured by the TIMING platform can distinguish the behavior of T cells before, during and after synapse formation, including motility, morphology, cytokine secretion, time to form synapse, duration of synapse, and time from synapse formation until death occurs in the target cell. Further, the TIMING platform can use these parameters to identify unique groups of cells and distinguish various cell behavior upon cell activation. Artificial intelligence algorithms enable high-throughput measurements of properties from single cells and cell-cell interactions which cannot be assessed with image acquisition over a bulk population of T cells and target cells (**Figure 1A**). TIMING was used to characterize the anti-viral activity of T cells obtained from a convalescent donor. Immortalized cells pulsed with peptides from spike (S) protein were used as targets to mimic infection by SARS-CoV-2. TIMING was adjusted to image sets of single T cells at short intervals of five minutes to quantify the behavior and movement of cells along with their cell-cell interactions in the presence of stimulator cells.

**Figure 1:**
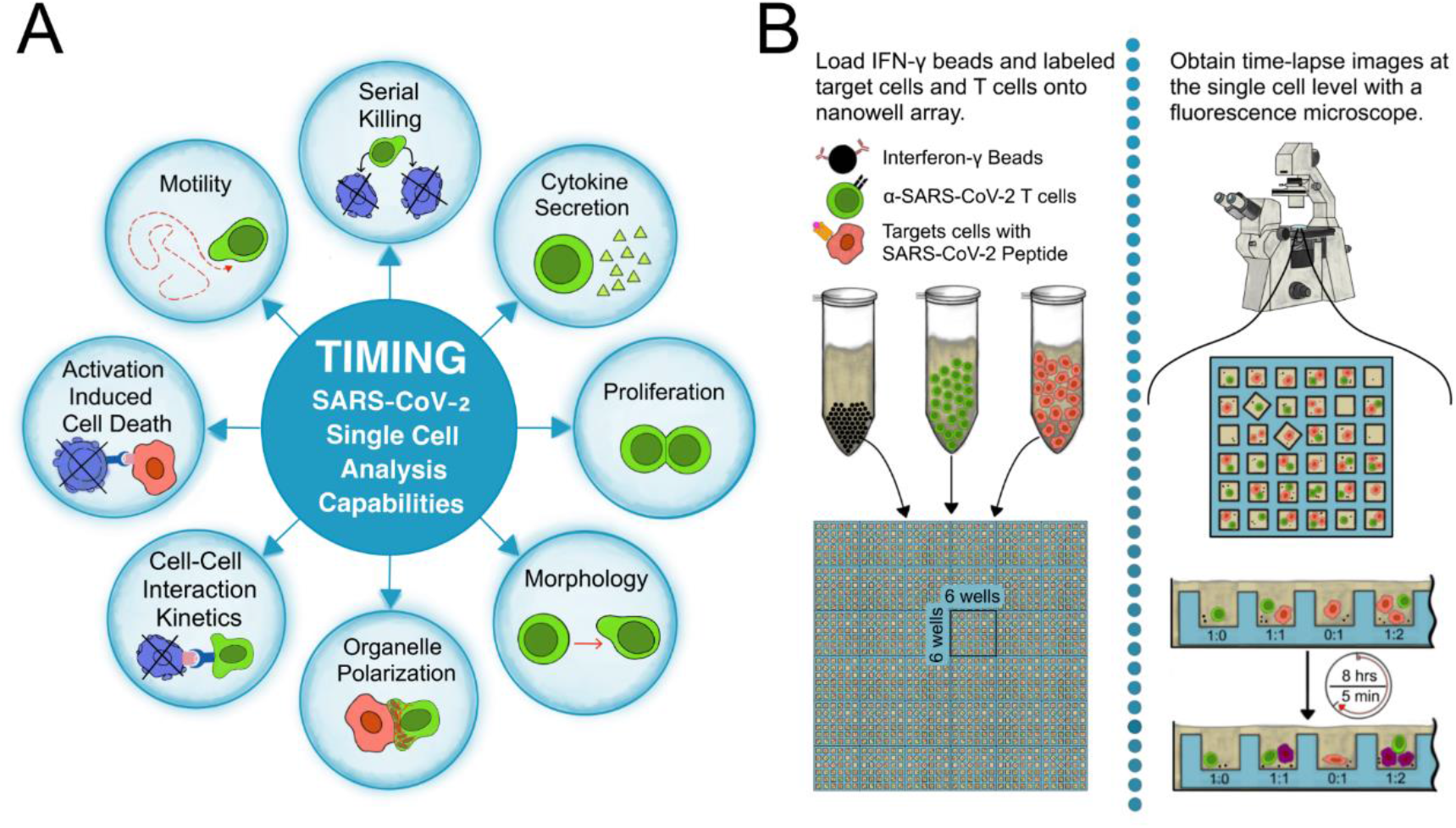
(**A**) Parameters measured by the TIMING platform. (**B**) The workflow of a TIMING assay. This platform is available through fee-for-service at CellChorus (https://cellchorus.com).

The TIMING platform was able to visualize, monitor and analyze the dynamic properties of SARS-CoV-2 specific T cells in stimulator cells pulsed with peptides derived from SARS-CoV-2 S-protein. This revealed heterogeneity of T cells in their ability to mediate a suite of effector functions and in particular cytotoxicity and cytokine production. These data inform on the immune response of a patient with COVID-19 highlighting the complexity of T-cell responses.

## Material and Methods

SARS-CoV-2-specific HLA A2-restricted cytotoxic T cells (Cellero, Cat #1142) were cultured in RPMI 1640 supplemented with 10% FBS, 1% sodium pyruvate, 1% L-glutamine, 1% Pen/Strep, 1% HEPES, and 50 IU/mL IL-2. A375 cells (ATCC, Cat #CRL-1619) were cultured in Dulbecco’s Modified Eagle Medium (DMEM) supplemented with 10% FBS, 1% sodium pyruvate, 1% L-glutamine, 1% Pen/Strep, and 1% HEPES. A375 cells were confirmed for absence of mycoplasma contamination using real-time PCR. We used TIMING to evaluate T cell-mediated cytotoxicity at a single-cell scale as previously described (**Figure 1B**)^8^.

We prepared the PLL-g-PEG (SuSoS, Dübendorf, Switzerland) solution by dissolving 1mg of it in 10 mM HEPES buffer. We used PKH26 and PKH67 dye (Sigma-Aldrich, Saint Louis, MO, USA, Cat #PKH26GL and #PKH67GL) for A375 and T cells staining, accordingly. To prepare the capture beads for IFNγ detection, we coated the goat anti-mouse IgG Dynabeads (Thermofisher, Waltham, Mass, USA, Cat #11033) with 40 μg/mL anti-human IFNγ mAb 1-D1K (MABTECH, Stockholm, Sweden, Cat #3420-3-250) for 30 minutes followed by two washes. We used AF647-Annexin V (ThermoFisher, Waltham, Mass, USA, Cat #A23204) as a death marker and anti-CD28 antibody (ThermoFisher, Waltham, Mass, USA, Cat #16-0289-81, Clone CD28.2) to stimulate the secretion of IFNγ. We acquired all images using a Carl Zeiss Axio Observer (Jena, Germany) fitted with a Hamamatsu Orca-Flash sCMOS camera and a 20 × 0.8 NA objective. We collected and processed images using an in-house algorithm for tracking and segmentation of cells^7^. The steps to undertake TIMING are described below.

1. Oxidize the nanowell array (144 blocks, each with 36 individual nanowells) with air plasma and incubate with PLL-g-PEG solution for 20 minutes at 37°C.
2. Load the stained A375 target cells at density of 1 million/mL on the nanowell array and pre-incubated them with 5 μg/mL KLPDDFTGCV peptide (JPT, Germany) for 60 minutes.
3. Load the stained T cells and coated Dynabeads™ at 10 μg/mL on the nanowell array.
4. Cover the nanoarray with media containing AF647-Annexin V and 5 μg/mL anti-CD28 antibody.
5. Acquire images of the nanoarray for 8 hours at 5 minutes intervals using an epifluorescent microscope.
6. Incubate the array with 10 μg/mL biotinylated anti-human IFNγ 7-B6-1 (MABTECH AB, Stockholm, Sweden, Cat #3420-6-250) and then 12 μg/mL AF647-Streptavidin (Jackson ImmunoResearch, West Grove, Penn., USA, Cat #016-600-084) for 30 minutes.
7. Acquire images of the array for IFNγ detection using an epifluorescent microscope (**Figure 1B**).

## Results

### Individual CTLs exhibit heterogeneity in killing efficacy

We deposited individual cells from a clonal human CD8^+^ cytotoxic T lymphocytes (CTL) population specific for the SARS-CoV-2 spike epitope ^424^KLPDDFTGCV^433^ (hereafter, DD9), restricted by HLA-A*02, into the wells of a nanowell array. We performed an alignment of the S protein across the major circulating SARS-CoV-2 variants that showed the DD9 epitope is conserved across all major variants including the Omicron variant (**Supplementary Figure S1**). As targets, we used the HLA-A*02 cell line, A375 cells, pulsed with the DD9 peptide. This process of sequential loading of cells yielded 308 nanowells containing a single effector (E) and one or two targets (T). Within the same nanowell array, 687 nanowells had only a single target cell and 659 nanowells had only a single effector cell. These served as control experiments for monitoring spontaneous target and effector apoptosis (**Supplementary Figure S2A**). We subsequently loaded the nanowells array with beads functionalized with antibodies against interferon gamma (IFNγ). The beads were used to track IFNγ secretion at the single-cell level and map IFNγ secretion onto the multiple dynamic parameters measured using microscopy.

Tracking the interaction between effector and target cells revealed marked heterogeneity between individual CTLs. We observed that within all nanowells bearing 1E:1T (N_total_ = 270), 36% of the CTLs formed a stable synapse (conjugation of >= 10 min) [**Figure 1A**]. Even among CTLs that contributed to such a synapse, only 27% of these cells induced apoptosis within the target cells (**Figure 2A, Supplementary Figure S2B**). These results demonstrate that there is intra-clonal functional heterogeneity within CTLs and that a minority of effectors contribute to the majority of killing observed. These results cannot be obtained when evaluating effector function across populations (**Supplementary Figure S3**).

**Figure 2:**
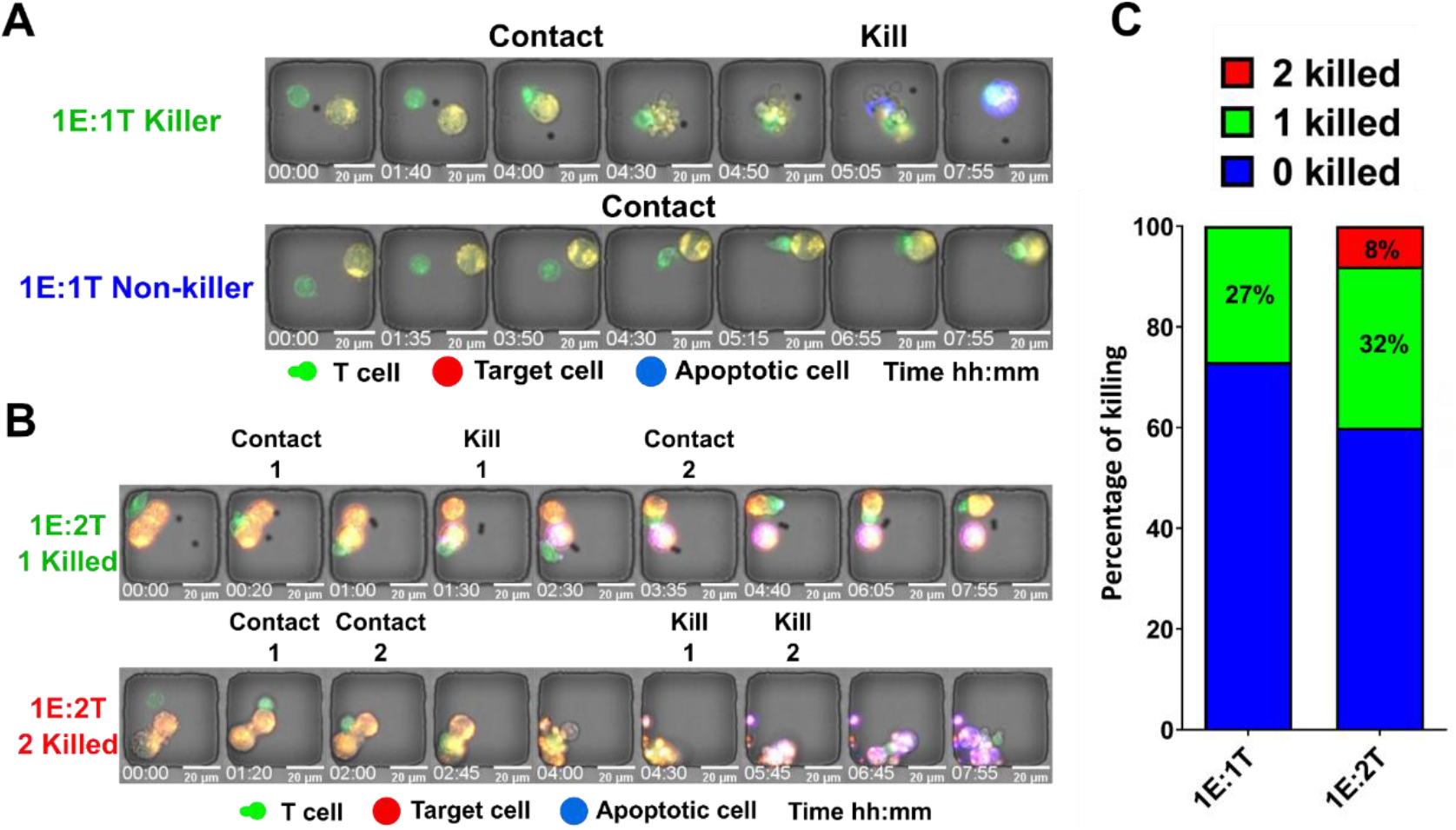
Cytotoxic T cells specific for SARS-CoV-2 spike protein epitope are functionally heterogeneous. (**A**) The micrographs of a killer and a non-killer T cell after contacting target cells (1E:1T). (**B**) The micrographs of a killer and a serial killer T cell (1E:2T). (**C**) Percentage of killer T cells in two E:T ratios (1E:1T, 1E:2T). The ‘2 killed’ subpopulation indicates serial killer T cells.

The superior killing ability observed by a sub-population of CTLs can be attributed to a subpopulation of serial killer T cells. TIMING allows the direct identification and quantification of serial killing by individual T cells. This was achieved by analyzing nanowells bearing 1E:2T (**Figure 2B, Supplementary Video S1**). In these nanowells, 32% of T cells lysed one target whereas 8% of CTLs participated in serial killing (eliminated both targets) [**Figure 2C**]. Collectively, these data demonstrate that most of the killing capacity of the virus-specific T cells is from a small fraction of killer and serial killer effector cells.

### Kinetics of T-cell mediated killing

The formation and duration of a synapse between T cells and targets are critical parameters that describe the efficiency with which killing occurs. The kinetics of T cell and target interactions were segmented into three defined stages: tSeek, the amount of time it takes T cells to locate targets and establish a stable synapse (contact lasting at least 10 min); tContact, the duration of stable synapse; and tDeath, the time from establishing a stable synapse until induction of apoptosis in a target cell (**Figure 3A**). tSeek was shorter in killer T cells in comparison to non-killer T cells that formed a synapse but did not induce apoptosis in any target cells (**Figure 3B,E, Supplementary Video S2**). After the establishment of the synapse, killer T cells delivered the lytic hit and detached prior to the induction of apoptosis in a target cells (**Figure 3C**). In contrast to killer T cells, non-killer T cells failed to efficiently terminate the synapse and remained conjugated to the target cells significantly longer (**Figure 3D**). Collectively, these results suggest that killer T cells are time efficient at the establishment of synapse, termination of synapse, and subsequent detachment. It has been previously demonstrated that this cycle of efficient conjugation and detachment is associated with sustained killing in lymphocytes^9^.

**Figure 3:**
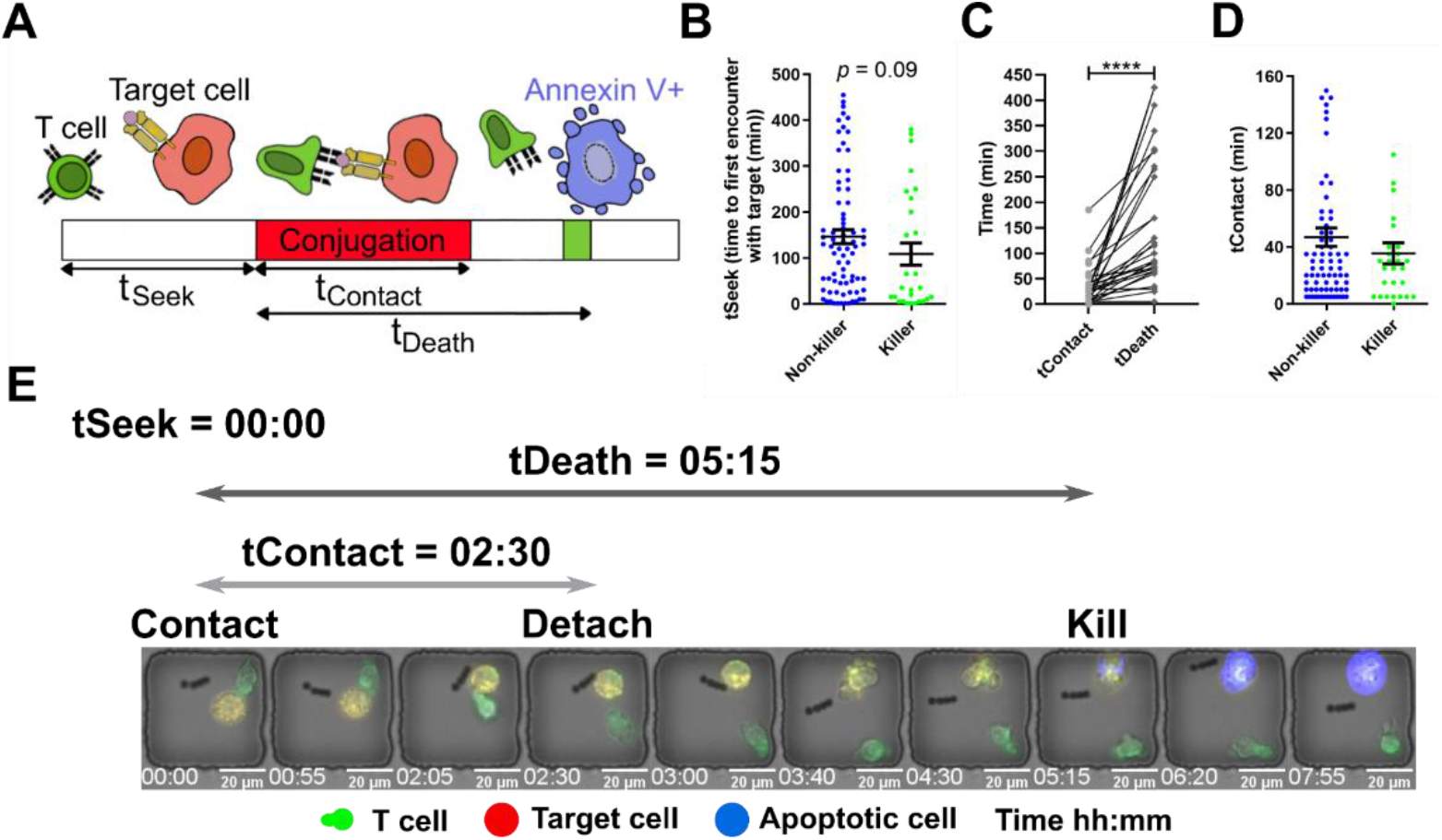
Kinetics of CTL killing. (**A**) Schematic of the dynamic parameters measured by the TIMING platform. (**B**) Time to first contact with target (tSeek) between killer and synaptic cells with mean ± SEM and Mann Whitney t test. (**C**) Comparison of contact duration (tContact) and time to induce target death (tDeath) for single cells with Mann Whitney t test and **** *p* < 0.00001. (**D**) Duration of stable synapse (tContact) between killer and synaptic cells with mean ± SEM. (**E**) A micrograph of a killer T cell detaching from a target cell before the induction of apoptosis.

### Quantifying the motility of single cells

A key property of T cells is their ability to migrate in search of target cells. We tracked the motility of individual T cells in nanowells populated with 1E:0T (N_total_ = 659) and observed that there was a gradient in effector cell motility ranging from 0.1 to 3.0 μm/min. We also evaluated the dynamic shape of T cells and observed that migrating cells had a lower aspect ratio of polarization in comparison to non-migrating T cells, which were predominantly circular (**Figure 4A**). We observed significant differences in the frequency of synapse forming T cells based on their motility (**Figure 4B, Supplementary Video S3**). These observations highlight the fact that migration and seeking of the target cells is a pre-requisite for T cell effector functionality such as cytotoxicity.

**Figure 4:**
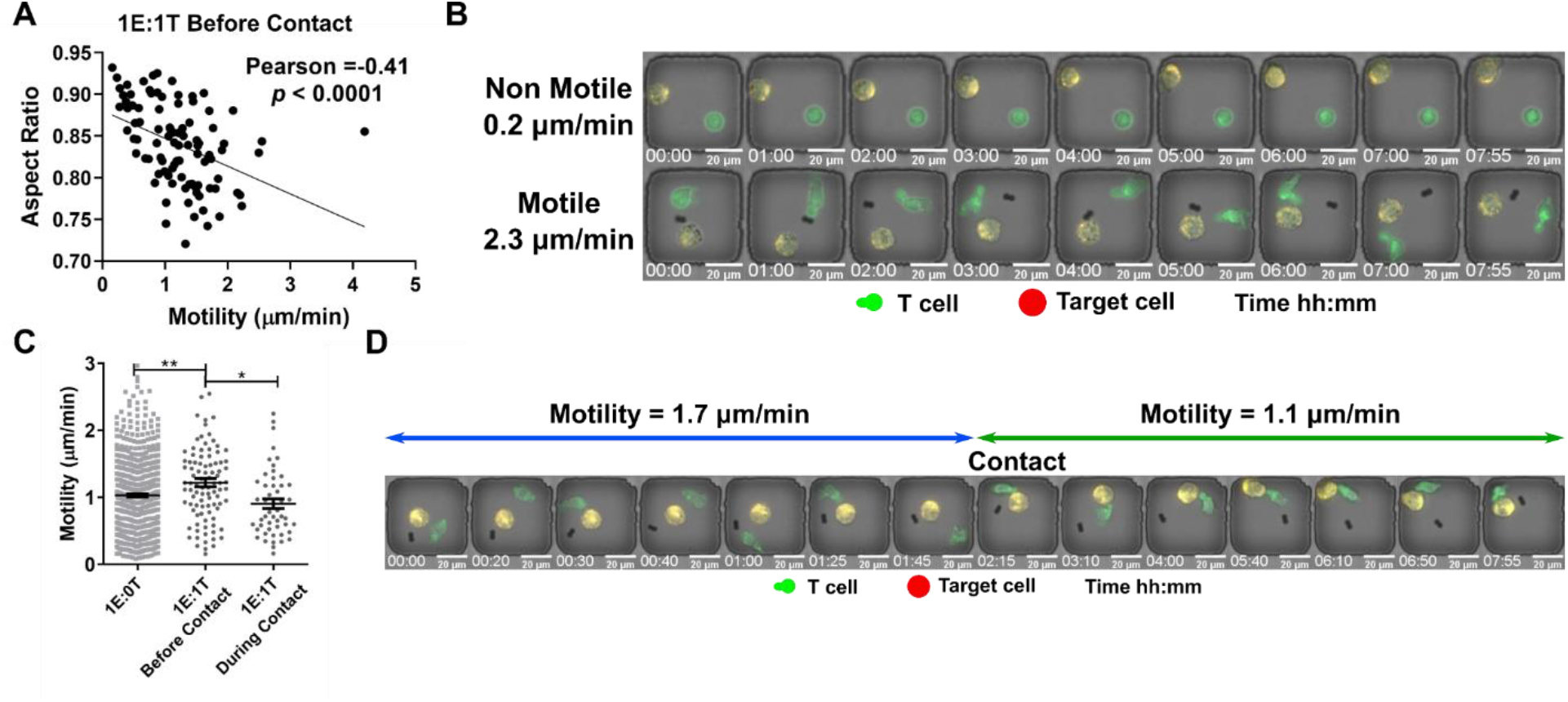
The motility and polarization of CTLs. (**A**) The negative correlation of aspect ratio and motility of T cells before contacting a target cell (1E:1T). (**B**) The micrographs of a non-motile and a motile T cell before contacting a target cell (1E:1T). (**C**) The motility of T cells in absence (1E:0T) and presence of a target cell (1E:1T). In cases of 1E:1T, the motility before and during contact is presented. Each data point represents a single cell with mean ± SEM. Mann Whitney t test with * *p* < 0.01, ** *p* < 0.001. (**D**) The micrograph of a T cell before and during contacting a target cell.

We tracked the motility of individual T cells in nanowells in two states: with and without conjugation to a target cell. The motility of T cells without conjugation to a target cell (1E:1T) increased compared to T cells located in wells without a target cell (1E:0T). This suggests that the T cell motility is activated by co-incubation with target cells. Upon conjugation to a target cell, however, there was a noticeable drop in the motility of individual T cells, consistent with the known immunobiology of T cells in arresting motility to optimize effector function (**Figure 4C and D)**^10^.

### Mapping the secretion of killing activity and cytokine production at the resolution of individual T cells

In addition to killing, CTLs are capable of secreting antiviral cytokines such as IFNγ. We have previously shown that killing and cytokine secretion can be decoupled within virus-specific T cells^11^. To investigate the relationship between killing and cytokine secretion at single-cell resolution, we tracked individual T cells that formed a synapse (1E:1T) and mapped the polyfunctionality of killing activity and cytokine production. Polyfunctional cells comprised only 11% of the population whereas the dominant population of T cells were non-functional (Killing^neg^IFNγ^neg^) (**Figure 5A, Supplementary Video S4**). Next, we compared the rate of secretion of IFNγ between killer and non-killer T cells (based on intensity of immunofluorescence on the bead^12^) and observed that killer CTLs secreted significantly higher amounts of IFNγ compared to non-killer T cells (**Figure 5B**). Collectively, these results illustrate that an elite subpopulation of T cells is capable of both killing and IFNγ secretion.

**Figure 5:**
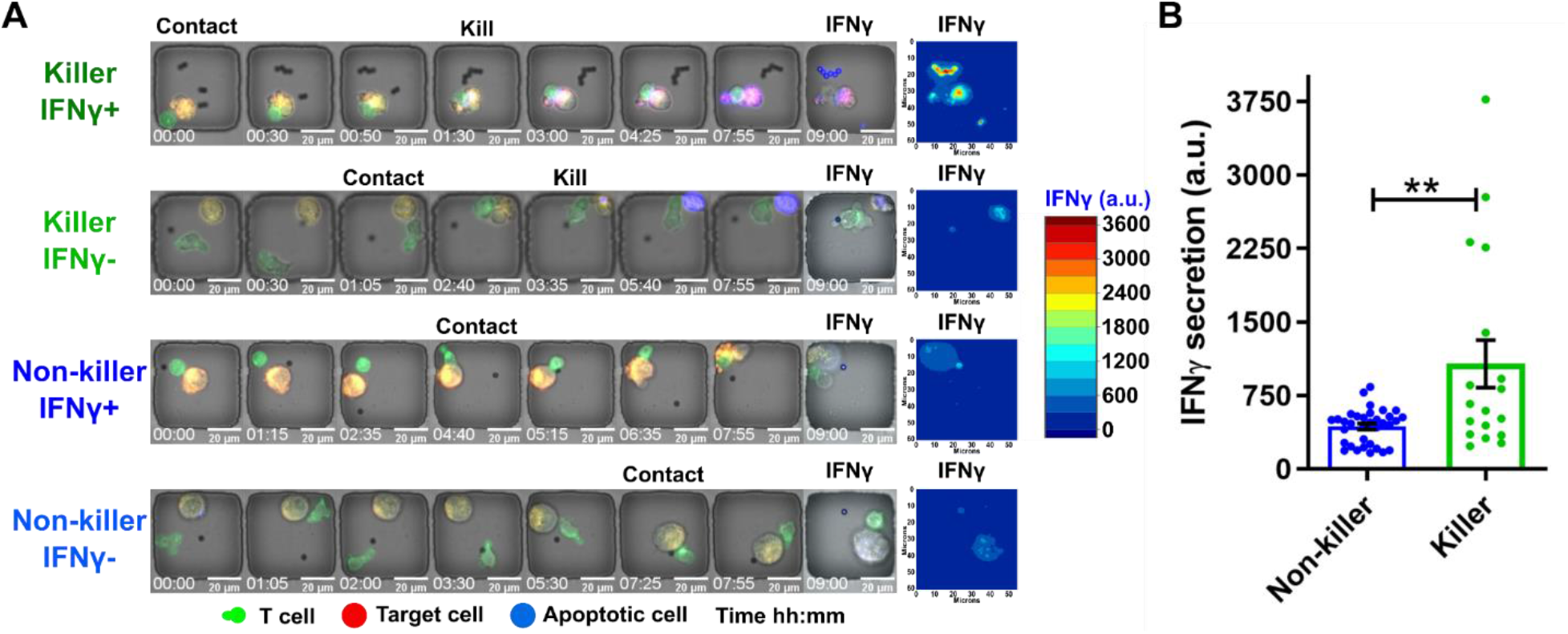
Quantification of IFNγ secretion from CTLs. (**A**) The micrographs of two killer T cells and two non-killer T cells with high and low secretion of IFNγ (all 1E:1T). The intensity of detection antibody on the bead (specific for IFNγ) at end point images represents the secretion of IFNγ. The contour map shows the intensity of IFNγ on the beads. (**B**) The quantified secretion of IFNγ from each cell (killer and non-killer). Each data point represents a single cell with mean ± SEM, Mann Whitney t test with ** *p* < 0.001.

### Serial killer CTLs have reduced activation-induced cell death occurrences

Activation-induced cell death (AICD) is a negative regulator of T cells and can titrate the potency and efficacy of effector cells^13^. Since TIMING serially tracks apoptosis occurring in both T cells and targets, we quantified the frequency of AICD in T cells as a function of their killing capacity (**Figure 6A, Supplementary Video S5**). Within all nanowells with evidence of T cell synapse formation (1E:1T), we observed an elevated frequency of killer CTLs undergoing AICD compared to non-killer CTLs. This observation was reversed when comparing serial killer CTLs (1E:2T) to non-killer CTLs (1E:2T) [**Figure 6B**]. These observations revealed that serial killer CTLs are more resistant to AICD than non-killer CTLs.

**Figure 6:**
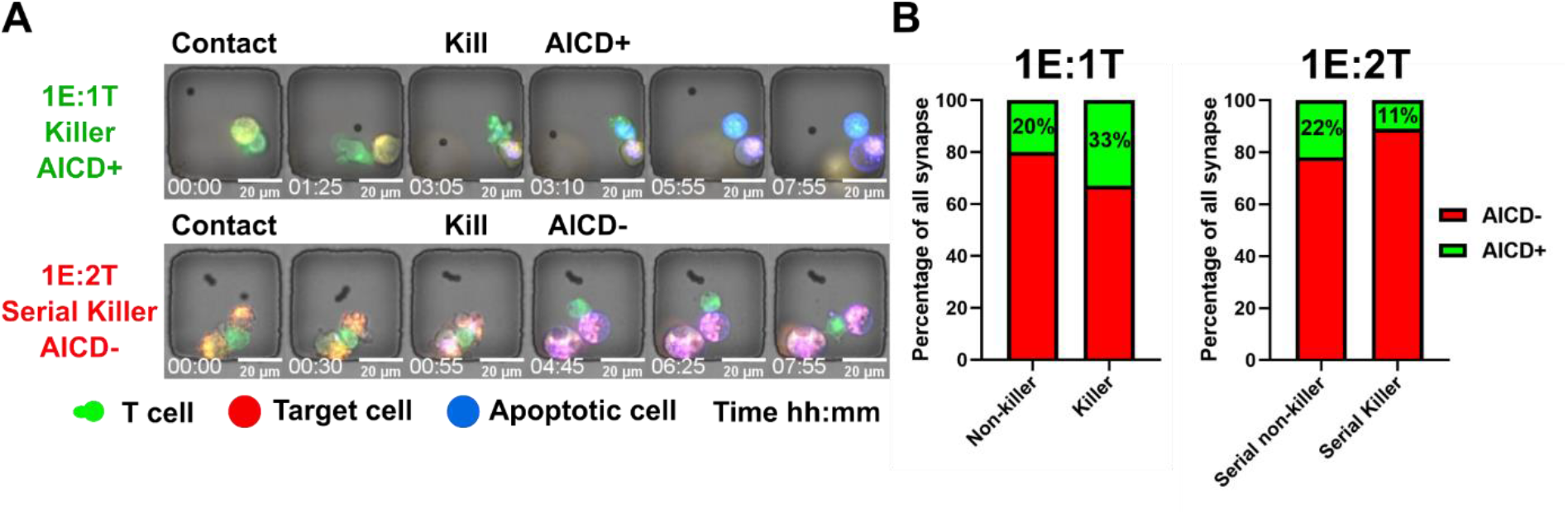
Quantifying AICD in T cells. (**A**) Micrographs of a killer T cell (1E:1T) undergoing AICD and a serial killer T cell (1E:2T) resistant to AICD. (**B**) Percentage of killers and serial killer T cells that either survived or underwent AICD.

## Conclusion

Current commercial vaccines for SARS-CoV-2 significantly reduce mortality and morbidity, but the rise of new mutations may challenge these successes^14^. Understanding a complete immune response by evaluating cellular immunity as well as humoral response is critical to appreciate the potential for vaccines to control new viral variants and to help understand those who have recovered from viral-induced disease. Variants of SARS-CoV-2 began to emerge towards the end of 2021 and raised the possibility that these viruses may evolve to elude immune defenses after infection or vaccination^14^. Studies have since investigated whether immunity is protective by examining responses in people who had contracted and recovered from SARS-CoV-2 prior to the widespread emergence of variants. These data reveal that CD8^+^ T cells remain active against the virus^4,15^.

The TIMING platform provides a new approach to understanding the complexity of the cellular immune response. This microscopy technique offers multiple advantages to dissecting T-cell responses to SARS-CoV-2. First, it synchronously monitors thousands of individual T cells as they move in their environment. Second, it analyzes T cells that recognize target cells bearing virally derived peptides. Third, it reveals T cells that are capable of polyfunctionality based on killing, serial killing and secretion of IFNγ. These three groups of data are integrated to reveal the dynamic interplay between T cells and target cells. To the best of our knowledge, this is the first report directly demonstrating that a subpopulation of CTLs specific for a conserved SARS-CoV-2 epitope are serial killers. The ability of CTLs to efficiently kill targets presenting conserved epitopes illustrates the power of T-cell mediated immunity and emphasizes the need to study cellular immune responses in the context of infection and vaccines.

More broadly, TIMING is an integrated platform that leverages advances in microfabrication, microscopy and AI to enable the profiling of thousands of timelapse microscopy experiments in parallel. As we have shown previously, the ability to integrate the dynamic profile with downstream clonal expansion or linked transcriptional profiling makes TIMING the only platform capable of generating dynamic multi-omic datasets^16,17^.

## Acknowledgements

We would like to thank Dr. Melisa Martinez Paniagua and Mel Montalvo for their help on the Cytation bulk cytotoxicity assay. We would like to thank Ali Rezvan for the fabrication of the nanowell arrays.

## Author Contributions

MF performed experiments.

MF, LC, NV, and DM interpreted and analyzed data.

LC, LJNC, NV, and DM provided advice on experiments and commented on the manuscript.

MF, NV, and DM designed and directed the study.

MF, LC, NV, and DM wrote the manuscript.

## Supplemental Materials

Additional figures.

Supplementary videos.

## Financial disclosure

NV was supported by the NIH (R01GM143243), CPRIT (RP180466), MRA Established Investigator Award (509800), NSF (1705464), CDMRP (CA160591), and Owens foundation.

DM, NV, MF, LC and LJNC have equity ownership in CellChorus.

LJNC receives royalties from Bristol Myers Squibb (via City of Hope National Medical Center) and Immatics (via MD Anderson Cancer Center), holds receipts of intellectual property rights from Sangamo BioSciences, MD Anderson Cancer Center, and Ziopharm Oncology, and also has equity ownership interest in Targazyme, Ziopharm Oncology, Immatics, CellChorus, Secure Transfusion Services, AuraVax Therapeutics, and IterateBio. NV has equity ownership in AuraVax Therapeutics and owns intellectual property rights from UH.

## References

1. Steensels D, Pierlet N, Penders J, Mesotten D, Heylen L. Comparison of SARS-CoV2 Antibody Response Following Vaccination with BNT162b2 and mRNA-1273. JAMA 326(15), 1533–1535 (2021).

2. Zhong D, Xiao S, Debes AK, Egbert ER, Caturegli P. Durability of Antibody Levels After Vaccination With mRNA SARS-CoV-2 Vaccine in Individuals With or Without Prior Infection. JAMA 326(24), 2524–2526 (2021).

3. Lee S, Miller SA, Wright DW, Rock MT, Crowe Jr JE. Tissue-specific regulation of CD8+ T-lymphocyte immunodominance in respiratory syncytial virus infection. Journal of virology 81(5), 2349–2358 (2007).

4. Redd AD, Nardin A, Kared H, et al. CD8+ T cell responses in COVID-19 convalescent individuals target conserved epitopes from multiple prominent SARS-CoV-2 circulating variants. medRxiv (2021).

5. Tan AT, Lim JM, Le Bert N, et al. Rapid measurement of SARS-CoV-2 spike T cells in whole blood from vaccinated and naturally infected individuals. The Journal of clinical investigation 131(17) (2021).

6. An X & Varadarajan N. Single-cell technologies for profiling T cells to enable monitoring of immunotherapies. Current opinion in chemical engineering 19, 142–152 (2018).

7. Liadi I, Singh H, Romain G, et al. Individual motile CD4+ T cells can participate in efficient multikilling through conjugation to multiple tumor cells. Cancer Immunology Research 3(5), 473–482 (2015).

8. Merouane A, Rey-Villamizar N, Lu Y, et al. Automated profiling of individual cell–cell interactions from high-throughput time-lapse imaging microscopy in nanowell grids (TIMING). Bioinformatics 31(19), 3189–3197 (2015).

9. Netter P, Anft M, Watzl C. Termination of the activating NK cell immunological synapse is an active and regulated process. The Journal of Immunology 199(7), 2528–2535 (2017).

10. Deguine J, Breart B, Lemaître F, Di Santo JP, Bousso P. Intravital imaging reveals distinct dynamics for natural killer and CD8(+) T cells during tumor regression. Immunity 33(4):632–44.

11. Varadarajan N, Julg B, Yamanaka YJ, et al. A high-throughput single-cell analysis of human CD8+ T cell functions reveals discordance for cytokine secretion and cytolysis. The Journal of clinical investigation 121(11) (2011).

12. An X, Sendra VG, Liadi I, et al. Single-cell profiling of dynamic cytokine secretion and the phenotype of immune cells. PLoS One 12(8), e0181904 (2017).

13. Zhong X, Matsushita M, Plotkin J, Riviere I, Sadelain M. Chimeric Antigen Receptors Combining 4-1BB and CD28 Signaling Domains Augment PI3kinase/AKT/Bcl-XL Activation and CD8+ T Cell-mediated Tumor Eradication. Molecular Therapy 18(2), 413–420 (2010).

14. Scott L, Hsiao N, Moyo S, et. al. Track Omicron’s spread with molecular data. Science 374(6574), 1454–1455 (2021).

15. Alter G, Yu J, Liu J, et al. Immunogenicity of Ad26. COV2. S vaccine against SARS-CoV-2 variants in humans. Nature 596(7871), 268–272 (2021).

16. Fathi M, Joseph R, Adolacion JRT, et al. Single-Cell Cloning of Breast Cancer Cells Secreting Specific Subsets of Extracellular Vesicles. Cancers (Basel) 13(17), 4397 (2021).

17. Bandey IN, Adolacion JRT, Romain G, et al (2021). Designed improvement to T-cell immunotherapy by multidimensional single cell profiling. Journal for ImmunoTherapy of Cancer 9(3).

