## Supplementary Materials for "Cytotoxic T lymphocytes targeting a conserved SARS-CoV-2 spike epitope are efficient serial killers"

**Supplementary Figure 1**

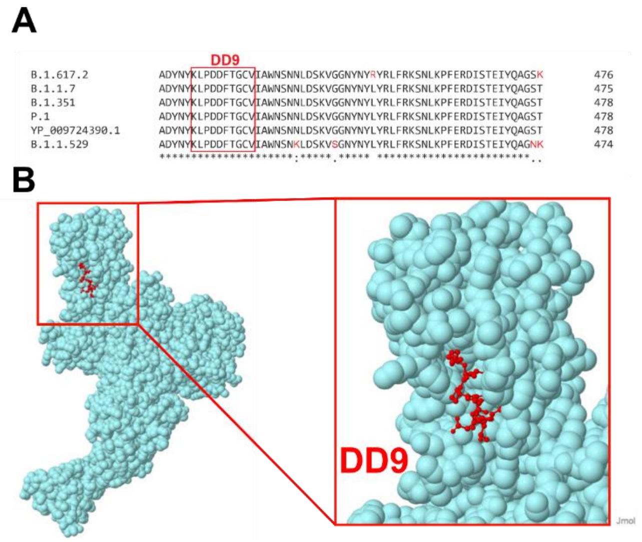


**Supplementary Figure 1:** (**A**) Multiple sequence alignment of the conserved epitope (DD9) on the spike protein of SARS-CoV-2 variants; YP_009724390 (Wuhan), B.1.617.2 (Delta), B.1.1.7 (Alpha), B.1.351 (Beta), P.1 (Gamma), and B.1.1.529 (Omicron). All sequences were downloaded from NCBI and aligned by Clustal Omega. (**B**) 3D structure of SARS-CoV-2 spike protein (PDB ID 6VYB) highlighted with DD9 peptide in red (Constructed in Jmol: an open-source Java viewer for chemical structures in 3D. http://www.jmol.org).

### Supplementary Figure 2

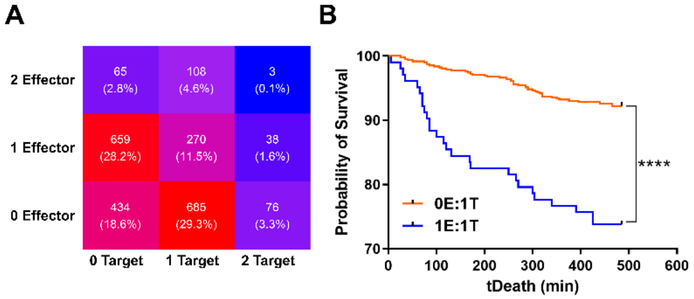


**Supplementary Figure 2:** (**A**) The distribution of T cells (effector) and target cells (target) in the nanowell array. (**B**) The Kaplan Meier survival curve of target cells in absence (0E:1T) and presence of one T cell (1E:1T).

**Supplementary Figure 3**

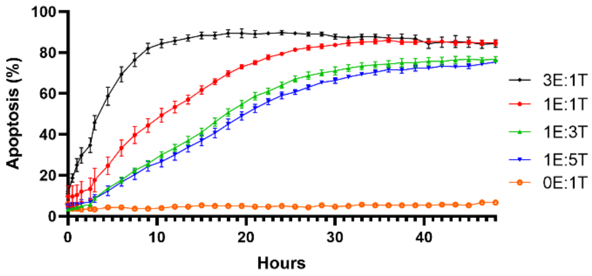


**Supplementary Figure 3:** The percentage of target cell apoptosis in different E:T ratios in a bulk assay for 48 hours (Cytation). All target cells were pre-incubated with peptide for 60 minutes before co-culturing.

**Supplementary Videos**

**Supplementary Video S1:** Representative TIMING Videos (each containing one T cell and one or more target cells) displaying killer, non-killer, and serial killer T cells.

**Supplementary Video S2:** Representative TIMING Videos displaying a killer T cell detaching from a target cell before induction of apoptosis.

**Supplementary Video S3:** Representative TIMING Videos displaying differences in motility and polarization of T cells.

**Supplementary Video S4:** Representative TIMING Videos displaying single nanowells containing killer or non-killer T cells with differences in IFNγ secretion.

**Supplementary Video S5:** Representative TIMING Videos displaying a killer T cell (1E:1T) succumbing to Activation-Induced Cell Death (AICD) and a serial killer T cell (1E:2T) resistant to AICD.
